# Climate change alters beneficial crop-microbe-invertebrate interactions

**DOI:** 10.1101/709089

**Authors:** Sharon E. Zytynska, Moritz Eicher, Michael Rothballer, Wolfgang W. Weisser

## Abstract

Increasing levels of CO_2_ and tropospheric ozone (O_3_) due to climate change are contributing to reduced plant health and unstable crop yield production^1^. The inoculation of plant roots with beneficial fungi or bacteria can increase plant health^2^. However, this is often studied under very controlled conditions and it is unknown how climate change or interactions with other species can alter the resulting benefits. Here we show that the rhizosphere bacterium *Acidovorax radicis* N35 can increase plant growth and reduce insect growth – with increased impact in a high-stress elevated O_3_ environment, but reduced impact under elevated CO_2_. In a fully-factorial climate chamber experiment we disentangled the impacts of climate factors (elevated CO_2_ and elevated O_3_) and biotic interactions (plant cultivar, sap-feeding insects and earthworms) on cereal growth and insect suppression mediated by *A. radicis* N35. Earthworms promoted plant aboveground growth, whereas *A. radicis* N35 promoted root growth, and overall plant growth was higher when both species were present. However, earthworms also promoted insect growth and therefore increased plant damage through herbivory. While *A. radicis* N35 inoculation was able to mitigate these negative effects to some extent under an ambient environment this was lost under climate change conditions. Our results show that knowledge-based solutions for sustainable agriculture should include biotic interactions and must be tested across variable climate change scenarios in order to build resilient cropping systems.

---

Crop protection is an important concern for future agriculture. The changing climate is reducing the predictability of crop yields^1,3^ and commonly-used insecticides are being banned for their potential role in insect decline^4,5^. Therefore, there is a need to develop knowledge-based solutions using technological advances as well as developing ecological-based approaches^6^. From an ecological standpoint, natural ecosystems are diverse with numerous species interacting to support a stable, well-functioning system. However, agricultural management practices can disrupt natural processes, leading to reduced ecosystem functioning^7^. While we cannot achieve sufficient crop yields through using only extensive farming, ecological intensification can be used to transfer our ecological knowledge of beneficial ecosystem functions into crop systems to our benefit^8,9^. One promising field of research is to manipulate the rhizosphere microbiome of plants^10^. Plants establish links with soil bacteria that can benefit plant growth through disease suppression, resistance to abiotic stress and nutrient acquisition^11^. Plant-associated bacteria and other beneficial soil organisms (e.g. ecosystem engineering earthworms) can even work together to further enhance plant health^12^. Creating soils that harbour organisms with complementary beneficial effects on the plant can mitigate losses in yield from reduced inputs^13^. As a consequence, using soil organisms as biostimulants is an active research field with great potential to achieve low-input agriculture^14,15^.

It is not clear, however, how plant-microbe interactions will effect plant growth under climate change. Increased frequency of short-term punctuated disturbances such as droughts, heat-waves and flooding, and longer-term increases in atmospheric carbon dioxide (CO_2_) and trophospheric ozone (O_3_) levels are impacting crop protection in various ways. Increases in CO_2_ tend to increase plant growth by enhancing their photosynthetic rate, but C3 plants (including cereal crops) can also experience a drop in nitrogen and trace elements potentially reducing the nutritional value for associated herbivores^16^. However, important crop pests such as phloem-feeding aphids tend to perform better under elevated CO_2_^17^ leading to more pest outbreaks in the future. An increase in tropospheric ozone levels is linked to by-products of fossil fuel combustion, with consequences for reduced plant yield, biomass, and increased susceptibility to pathogens^18^. However, ozone can also induce plant defence pathways, e.g. PR-proteins β-1,3-glucanases and chitinases^18^, that are involved in plant resistance to sap-feeding insects^19^. Therefore, we must understand outcomes of plant-pest interactions across variable environments to fully understand future implications and impacts on our food systems.

In a controlled fully-factorial experiment, we inoculated barley plant (*Hordeum vulgare* L.) seedlings with the rhizobacterium *Acidovorax radicis* N35^20^ (herewith, *A. radicis*). We assessed the effect of this on plant shoot and root growth and the growth rate of aphid insect pests across different biotic (barley cultivar, aphids, and earthworms) and climate environments (elevated CO_2_, elevated O_3_, and combined eCO_2_+eO_3_). We measured the initial viability of seedlings after transplantation (seedling growth from day 5 to day 8, before addition of aphids and earthworms), aboveground vegetative growth before tillering (plant growth from day 8 to day 22 after addition of aphids and earthworms), root growth (day 5 to day 22), and aphid number at day 22 (after 14 days’ growth) (Extended Data Fig. 1). Barley plant shoot and root length during the experimental period is a good predictor of dry biomass and final yield (Extended Data Fig. 2). The data were analysed using linear models on the full data (N=986; 4-11 replicates per treatment; Extended Data Fig. 1) and on paired data (N=474 pairs), which calculated the log-response ratio between plants inoculated with *A. radicis* to control plants within all other treatments. We calculated aphid density (number of aphids per cm of plant) to account for variable plant growth. The root-associated bacterial community (day 22 plants) was identified using 16S metabarcoding sequencing to analyse changes in diversity and composition across the treatments.

Our analyses uncovered multiple interactions between the climate and biotic factors on plant growth and aphid density, meaning that the effect of one factor depended on others (Extended Data Table 1). We also identified a number of general effects including increased plant growth under elevated CO_2_ and decreased growth under elevated O_3_ across all barley cultivars and other treatments^18^ (Fig. 1; Extended Data Table 1; Extended Data Fig. 3a-c). Earthworms increased plant growth, especially aboveground shoot growth (Fig. 1; Extended Data Fig. 3g-h), whereas the inoculated rhizobacterium *A. radicis* had a stronger growth promotion effect on the plant roots than on aboveground tissues (Fig. 2). The effect of inoculation by *A. radicis* was greatest on the aboveground plant shoot very early on when the seedling was establishing in the soil (Fig 2a), with no effect later on (Fig 2b) despite the strong belowground root growth promotion^21^ (Fig. 2c). One cultivar showed the opposite response during seedling establishment, with a negative response to *A. radicis* across all climate treatments (Extended Data Figure 4a-d); however, this cultivar responded most strongly in root growth (Extended Data Figure 4i-l) suggesting energy allocation variation across the cultivars studied.

**Figure 1.**
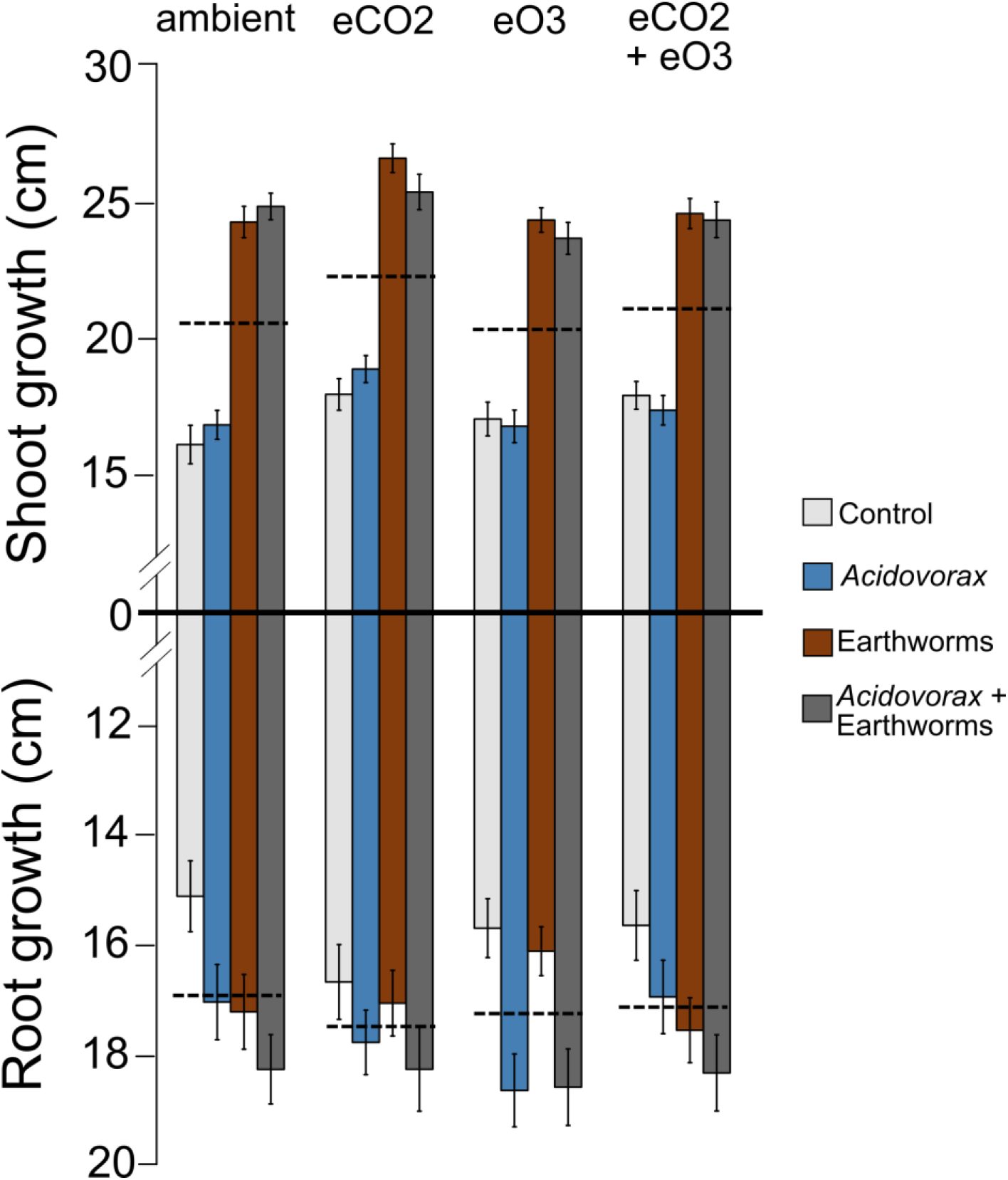
Plant shoot and root growth across biotic and climate treatments. The longer the combined root and shoot columns, the larger the overall growth of the plant. Dashed lines show the mean plant growth within each climate treatment averaged across all other treatment factors. Error bars are ±1 SE.

**Figure 2.**
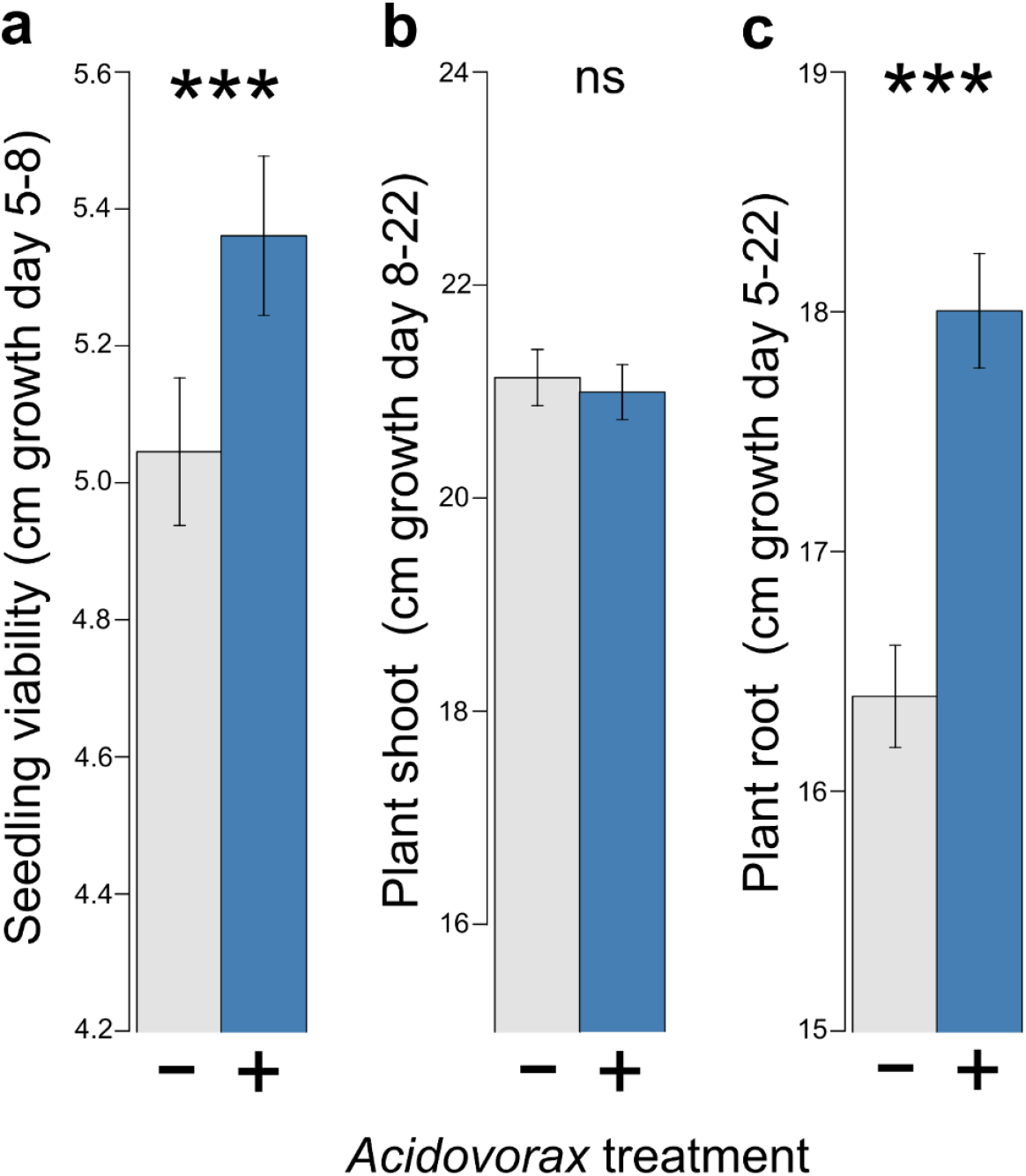
*Acidovorax radicis* inoculation on plant growth. (a) seedling viability, (b) plant shoot growth, and (c) plant root growth. The effect was strongest for root growth, and early seedling growth (seedling viability, for cultivars Barke, Grace and Scarlett), with no overall influence on later plant shoot growth. Error bars are ±1 SE. N=986

Elevated CO_2_ also increased aphid density (Extended Data Fig. 3d). Our measure of aphid density controlled for the increased plant growth in elevated CO_2_ and therefore this effect occurred through plant physiological changes^16^. Elevated CO_2_ can increase the relative abundance of essential amino-acids leading to increased aphid reproduction^22^, and influence plant defences against aphids through enhanced SA-signalling or suppression of JA- and ET-signalling pathways ^16^. Moreover, aphids release effector proteins that can alter local plant defences and manipulate nutrient availability, potentially allowing them to further benefit from elevated CO_2_^16^. Earthworms also increased aphid growth rates (Extended Data Fig. 3i). Meta-analyses have shown that earthworms can increase plant above- and belowground biomass (up to 30% in barley) predominantly through releasing nitrogen previously unavailable to the plant^23,24^. This may also benefit the aphids through increased amino-acid production and/or decreased plant defences^23,25^. In general, aphids reduced plant growth, likely through reducing the energy budget of the plant that would other be invested in growth as they feed on plant phloem-sap and induce plant defences^26^ (Extended data Fig. 3e-f).

While none of these main effects are unexpected, all treatment factors were also involved in higher-order interactions (Extended Data Table 1) and thus the individual impact of each must be considered in the wider context of the whole model ecosystem (context dependency). We found substantial variation across the four barley cultivars in their responses to the different environments (Extended Data Table 1), however this was predominantly through differences in the strength of effects (magnitude) rather than changes in the direction. This variation among cultivars can reduce the predictability of effects across a wider range of barley cultivars, yet we can harness this variation in future comparative analyses leading to a better understanding of the mechanisms underlying these interactions and benefiting future plant breeding.

To further understand how the benefits of *A. radicis* inoculation alter across the climate and biotic environments, we used matched pairs analysis comparing responses of control to treated plants (Extended Data Table 2). The response of the seedlings to the *A. radicis* inoculation depended on the climate environment, with reduced effect under elevated CO_2_ (Fig. 3a). There was also substantial variation across cultivars (Extended Data Fig. 4a-d). While Barke, Grace, and Scarlett cultivars responded overwhelmingly positively to *A. radicis* across the climate environments, Chevallier responded with reduced seedling viability. Grace showed highest beneficial effects of *A. radicis* under the eO_3_ stress environment and a negative response under elevated CO_2_ (Extended Data Fig. 4a-c). This may indicate increased potential of this plant cultivar to harness useful traits of beneficial organisms under stressed environments, but reduced ability in an optimal environment.

**Figure 3.**
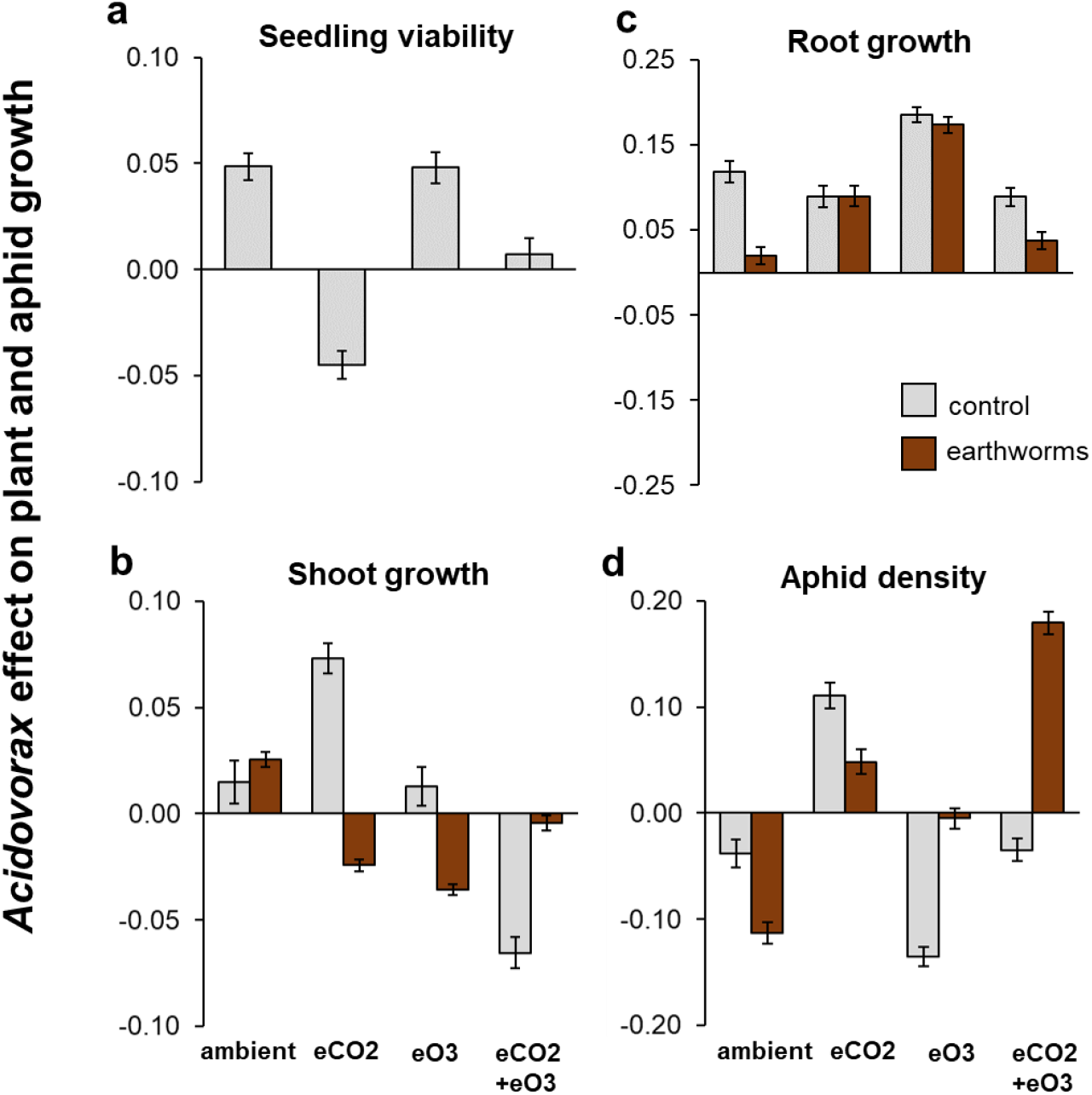
Effect of *A. radicis* inoculation from paired analysis. Data shows the log-response of plant and aphid growth traits comparing plants that were treated with A. radicis with one treated with a control solution. Error bars show the variance around this ratio.

The effect of inoculating plants with *A. radicis* on the later plant shoot growth was predominantly influenced by a combination of the earthworm biotic and climate treatments (earthworms x eCO_2_ x eO_3_ F_1,464_=5.39, P=0.021; Fig. 3b). Earthworms rarely enhanced any beneficial effects of *A. radicis* on later plant shoot growth, and even changed the direction from positive to negative in eCO_2_ or eO_3_ (Fig 3b). While *A. radicis* had much stronger beneficial effects on the plant root growth (Fig. 3c), even here the magnitude of these again varied across the biotic and climate environments (cultivar x earthworms x eCO_2_ x eO_3_ F_3,437_=3.90, P=0.009; Fig. 3c). Here, cultivars also showed substantial variation (Extended Data Fig. 4i-l). In the ambient environment, root growth of Barke showed a negative response to *A. radicis* yet this was reversed to be a relatively strong positive response under eCO_2_ and eO_3_, and especially when no earthworms were present (Extended Data Fig. 4i-l). Since both earthworms and *A. radicis* are beneficial for the plant, this suggests they do not facilitate one another to further enhance every plant growth trait but rather are complementary where one or the other provides the benefits. This results in overall greater total plant growth (shoot and root combined) when both beneficial species are present (t=6.27, df=3, P<0.001; Fig1, Extended Data Fig. 5).

For pest suppression, in an ambient environment *A. radicis* reduced aphid density to a greater extent when earthworms were present in the soil, e.g. in a biodiverse environment (Fig. 3d). This response changed under eCO_2_ (F_1,209_=3.98, P=0.047) when *A. radicis* was no longer able to supress aphids but rather increased them (Fig 3d); potentially through variation in plant nutrients or defence^16^. However, in the biodiverse environment with earthworms, this increase in aphid density from eCO_2_ and *A. radicis* was reduced. Under elevated O_3_ there was stronger suppression of aphid pests with no earthworms (F_1,209_=3.00, P=0.085; Fig 3d). This may be related to a trade-off in plant responses to beneficial earthworms and rhizobacteria under O_3_ stress^18^; however, there have been only a few studies looking at the direct impact of elevated ozone on soil organisms and thus these interactions need to be further investigated^27^. Under both eCO_2_+eO_3_ *A. radicis* could again supress the aphids but the presence of earthworms strongly altered this. Earthworm presence in agricultural fields depends on many factors including tillage practice, nitrogen fertilization and organic matter^28^, and therefore this context dependency is important to note when developing new methods for improving crop health.

In elevated CO_2_ environments, plants open their stomata for shorter periods of time and so it was expected that this would reduce negative impacts of elevated O_3_ in the combined environment^29^. We found that the results from the combined elevated CO_2_ and O_3_ treatment were not intermediate of the individual treatments and could not be predicted from them. This confirms a previous study that also found there was little association between the individual treatments and combined treatments^18^.

The microbial community analysis confirmed the presence of *A. radicis* in the rhizosphere at the end of the experiment, and also showed that increased abundance of *A. radicis* was correlated with increased plant growth and decreased aphid densities (Fig. 4a). The abundance of *Phenylobacterium* was also correlated with increased plant growth and decreased aphid densities, which could potentially be an alternative target for crop health. *Shinella* and *Porphyrobacter* bacteria were beneficial for both the plant and also the aphid, while others, such as *Burkholderia* were negative for the plant but somewhat positive for the aphids. *Burkholderia* is a known group containing many plant bacterial pathogens^30^, and subsequent analysis also showed that *A. radicis* inoculation and also earthworms reduced the abundance of these pathogenic bacteria (Extended Data Fig. 6) indicating one possible mechanism for the effects observed in our experiment. Overall, the inoculation of *A. radicis* did not alter the overall bacterial community on the barley roots (Fig. 4b). In contrast, aboveground aphid feeding (Fig. 4c) and the climate environment (Fig. 4d) significantly changed the root-associated microbiome.

**Figure 4.**
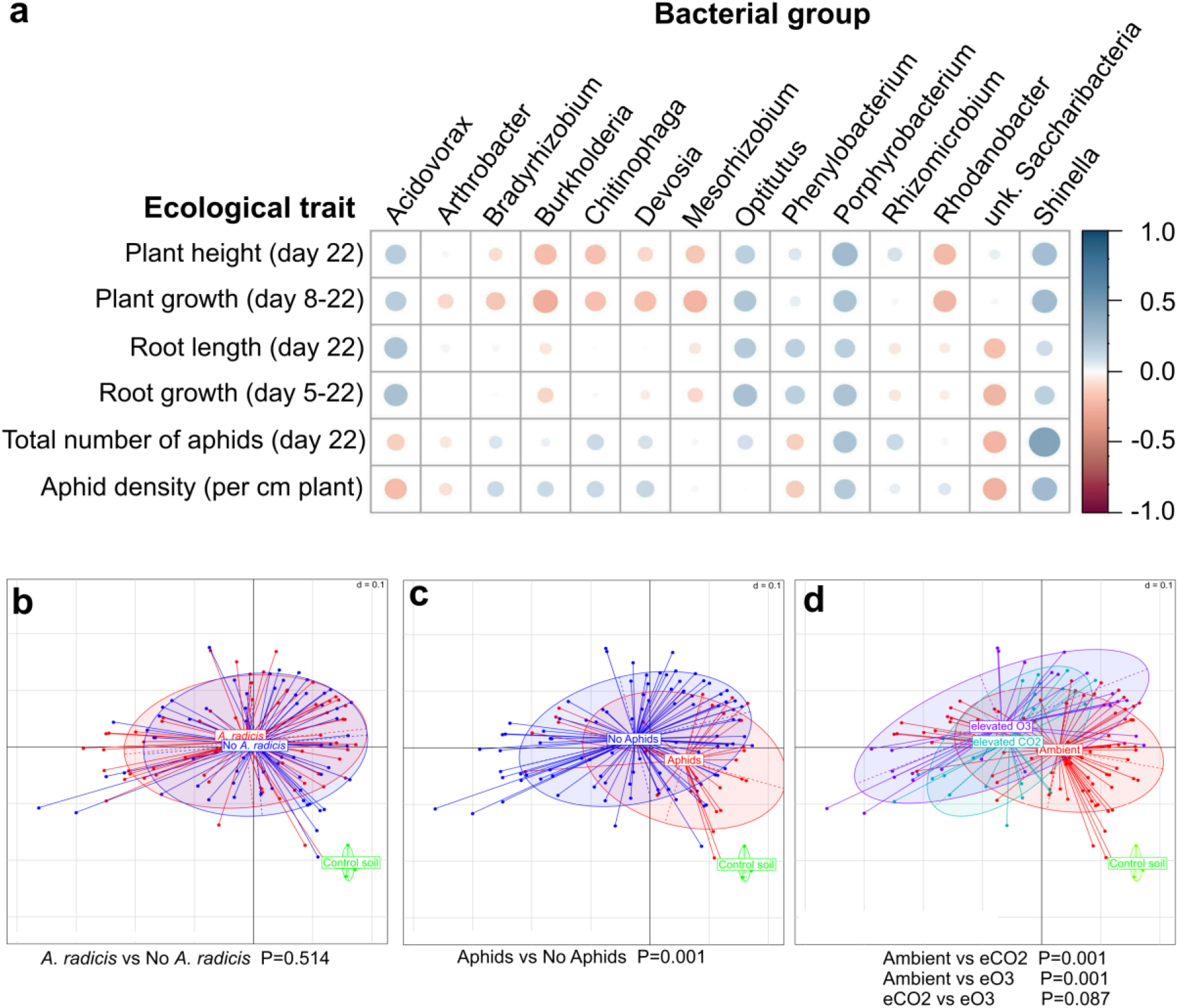
Statistical analyses of microbiome sequencing data with the Rhea pipeline. (a) Correlation plot showing negative (red) and positive (blue) correlations between different measured plant parameters and the abundance of detected genera. The bigger the circle the higher the significance (i.e. the lower the p-value), cutoff was set to p=0.05. (b-d) Multi-dimensional scaling plots of microbial profiles with d=0.1 meaning that the distance between two grid lines represents approximately 10% dissimilarity between the samples.

Our study showed that there is real promise for introducing beneficial soil species to benefit crop growth and reduce insect pests in sustainable agriculture. The context-dependency of the interactions was found to affect the strength of effects rather than the direction. This is important since complex interactions can lead to unpredictability in outcomes, yet we found that increasing complexity (diversity) of a system had overall beneficial outcomes. Under certain environments, only one beneficial species was required yet for many others the combination of beneficial species was a benefit for plant health. We highlight the need to include the effects of biotic and climate factors when developing knowledge-based ecological solutions in agriculture, and using soil organisms as biostimulants is a promising path towards achieving low-input agriculture^13–15^.

## METHODS

### Study system

Our study species included: (1) four European barley (*Hordeum vulgare*) plant cultivars: Barke (Saatzucht Breun GmbH), Chevallier (New Heritage Barley Ltd), Grace (Ackermann Saatzucht GmbH), and Scarlett (Saatzucht Breun GmbH); (2) the English grain aphid *Sitobion avenae* (L.) that had been maintained as low density stock populations on Barley cultivar ‘Kym’ in a climate cabinet for two years, the clone was originally from Goettingen University; (3) epigeic earthworms *Dendrobaena veneta* Rosa 1886, originally from wurmwelten.de and maintained in a Worm-Café^®^ for three years prior to the experiment; and (4) the rhizobacteria *Acidovorax radicis* N35 prepared by colleagues from the Helmholtz Zentrum Munich, along with a control solution containing no bacteria for seedling inoculation.

### Experimental design

The climate experimental treatments [elevated carbon dioxide, CO_2_ (+/−), elevated ozone, O_3_ (+/−)] were used at the level of an individual climate chamber with four chambers used: (1) ambient (~500 ppm day-time during high light periods, 600 ppm night-time); 0.02 ppb ozone), (2) elevated CO_2_ (700 ppm day-time during high light periods, 900 ppm night-time), (3) elevated O_3_ (constant 100 ppb), and (4) elevated CO_2_ and elevated O_3_. The experiment was run across three successive temporal blocks (runs), and chamber identity was changed across runs, such that each climate treatment was run in three different chambers across the experiment to avoid a chamber-treatment confounding effect.

The biotic experimental treatments [plant cultivar (Barke, Chevallier, Grace, Scarlett), *A. radicis* (+/−), earthworms (+/−), aphids (+/−)] were run at the level of an individual pot within a chamber. Within each run, three replicates of each biotic (plant cultivar, *A. radicis*, earthworm, aphid) treatment were made with each replicate allocated to one of three tables (randomised block design within run, within chamber). The total number of replicates in the design was nine, three per treatment per run.

The experimental design was fully-factorial, with three temporal blocks (runs) and blocks within chambers (tables). Table within chamber was not a significant block effect, indicating the high homogeneity of the climate chambers.

### Experimental set-up

Seeds were germinated between moistened filter paper for 5 days in the dark at room temperature. After this the seedlings were soaked in either *A. radicis*-containing solution or control solution for one hour. *A. radicis* was grown by inoculating the surface of NB plates, and incubated at 30 °C for 36 hrs. Then the cultures were resuspended in 10 mM MgCl_2_ with final suspension containing 10^9^ cells per ml. The control solution was 10 mM MgCl_2_, and 100μl Tween 20 was added to both bottles. Before transplantation, the length of the shoot and longest root of the seedlings was measured. Then, seedlings were planted into 10 cm pots (single seedling per pot) containing soil substrate (Floragard B Pot Medium-Coarse, pH 5.6, NPK 1-0.6-1.2) mixed with quartz sand at a 5:1 (soil:sand) ratio. Plants grew uncovered for three days, when shoot length (from top of the seed to the longest leaf) was again measured. Aphids were introduced to plants using a fine paintbrush to move two 4^th^ instar aphids from the stock populations (kept at low densities to avoid winged aphid production) onto the base of the plant shoot. From here, aphids will move up onto the plant where they feed, develop into adults and then begin to produce offspring within the next few days. Earthworms were first washed in tap water and placed into plastic tubs with moist tissue for 48 hours to remove gut contents. Then, five worms were introduced into the soils (at the same time as aphid infestation), with a total biomass 1.1-2.1g (biomass recorded).

All pots were covered with a 180 x 340 mm air-permeable cellophane cover (HJ Kopp GmbH, Germany) on the top, and organza mesh at the base of the pot, secured by two elastic bands. Plants were allowed to grow for 14 days under 20 °C, 65% RH (relative humidity), with 10 hours of full light (850 PAR), 8 hours of total darkness and a 3-hour sunrise/sunset gradient between these where light was gradually increased/decreased. At the end of the experiment, aphids were counted using hand tally-counters, ensuring a systematic method of counting each leaf from the base to the top. Plant shoot length (longest leaf) and root length (longest root) were measured, and earthworms extracted from the soil were washed, counted, and earthworm biomass measured. All five earthworms were recovered from 95.6% of pots, with only 2.5% of pots containing fewer than four earthworms (13/522 pots). Root material was collected and stored at −20 °C before DNA extraction for microbial community analysis.

### Phenotypic data analysis

Two approaches were used to analyse the phenotypic experimental data. All data were analysed in R 3.5.1 using RStudio (Version 1.1.463).

The first approach used standard linear models for variance partitioning of the data, where model response variables were (1) seedling viability: longest shoot length at day 8 minus longest shoot length at day 5 (cm), (2) plant growth: longest shoot length at day 22 minus longest shoot length at day 8 (cm), (3) Root growth: longest root length at day 22 minus longest root length at day 5, (4) aphid density: total number of aphids divided by the plant growth variable (day 22 – day 8) giving the number of aphids per cm of plant. All models included the experimental run as a blocking factor to control for variation across the three temporal blocks. Diagnostic plots of the models showed that standard linear models with a normal error distribution were suitable for all variables. Initial models included all main effects and interactions, and were simplified using a backwards stepwise method removing the least significant interaction terms one by one until a minimal adequate model is reached.

The second method focused on the effect of *A. radicis* inoculation on the same variables as above. However, here we used a matched pairs analysis that matched plants within treatments that had been inoculated with *A. radicis* compared to controls. We took care to only match plants from the same tables (achievable due to the randomised complete block experimental design used) to minimise differences due to variation within a chamber or across temporal runs. The absolute differences between these plants for each of the variables (seedling viability, plant growth, root growth, and aphid number) were then used to calculate the log-response ratio (lnRR, treated vs control). The lnRR values were then analysed using linear models using all main effects and interactions, thus determining the impact of these on the effect size (strength and direction) of *A. radicis* inoculation. Figures use the calculated mean effect size (lnRR) across treatment combinations and the associated variance (using the R package ‘metafor’).

### Microbial community barcoding

To assess the root-associated microbial community, 0.25-0.5 g of roots with attached soil was used for DNA extraction (Qiagen DNeasy PowerSoil Kit). The DNA extraction, amplification and sequencing was performed by AIM (Advanced Identification Methods GmbH, Munich). The V3-V4 region of the 16S rRNA gene was amplified using primers 341f (CCTACGGGNGGCWGCAG) and 785r (GACTACHVGGGTATCTAATCC), which showed the best coverage for bacteria and was most reproducible in a recent comparative evaluation^31^. A total of 7,382,326 paired end reads were recovered, with a median of 92.8% reads merged. Sequence data processing was performed using the IMNGS platform^32^ applying the UPARSE amplicon analysis pipeline^33^. Statistical evaluation was done with the Rhea pipeline for R^34^. The datasets supporting the conclusion of this article will be available through GenBank, the OTU table is available as a supplementary file.

## Supporting information

OTU Table microbial community

## Acknowledgements

The experiment was conducted within the climate controlled chambers of the TUMmesa (Technical University of Munich Model EcoSystem Analyser facility, Freising, Germany), and we thank R. Meier for maintenance and management of these chambers. We thank students R. Fahle, T. Braun, J. Kracht, and A. Janjic for help with data collection. Finally, we thank S. Meyer for comments that helped improve this manuscript.

## Author Contributions

SZ and WW designed the experiment, SZ and ME collected the experimental data. SZ analysed the experimental data and MR analysed the microbial community data. All authors interpreted the results, and SZ wrote the manuscript with all authors commenting.

## EXTENDED DATA

**Extended Data Figure 1.**
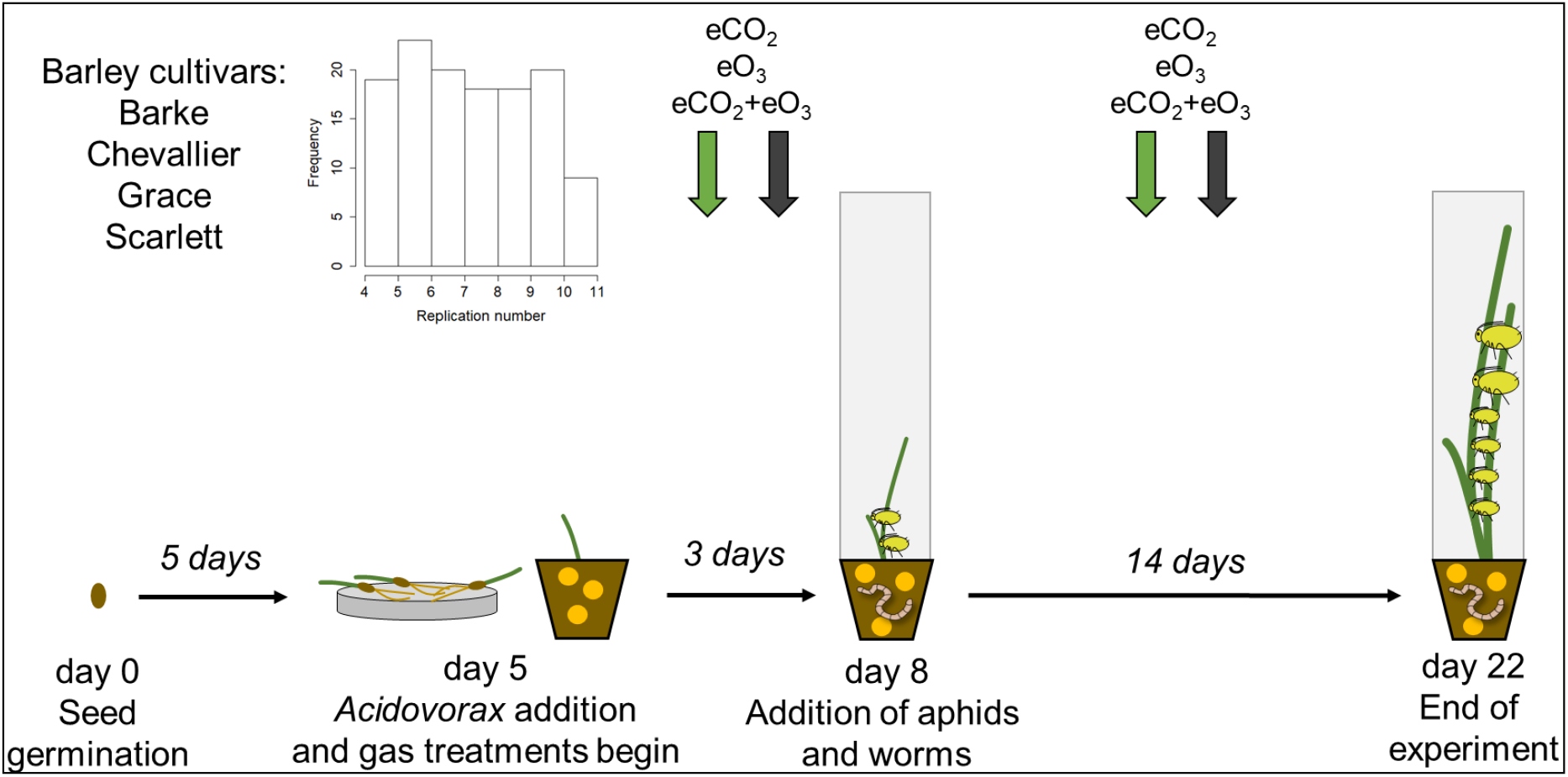
Summary of the experimental design. Inset show the distribution of the number of replicates for each treatment across the experiment, with a mean of 7.7, minimum of 4 and maximum of 11. Reasons for reduced replication are from low germination of one barley cultivar, and lack of growth of the barley after transplantation, which was not linked to any experimental treatment.

**Extended Data Figure 2:**
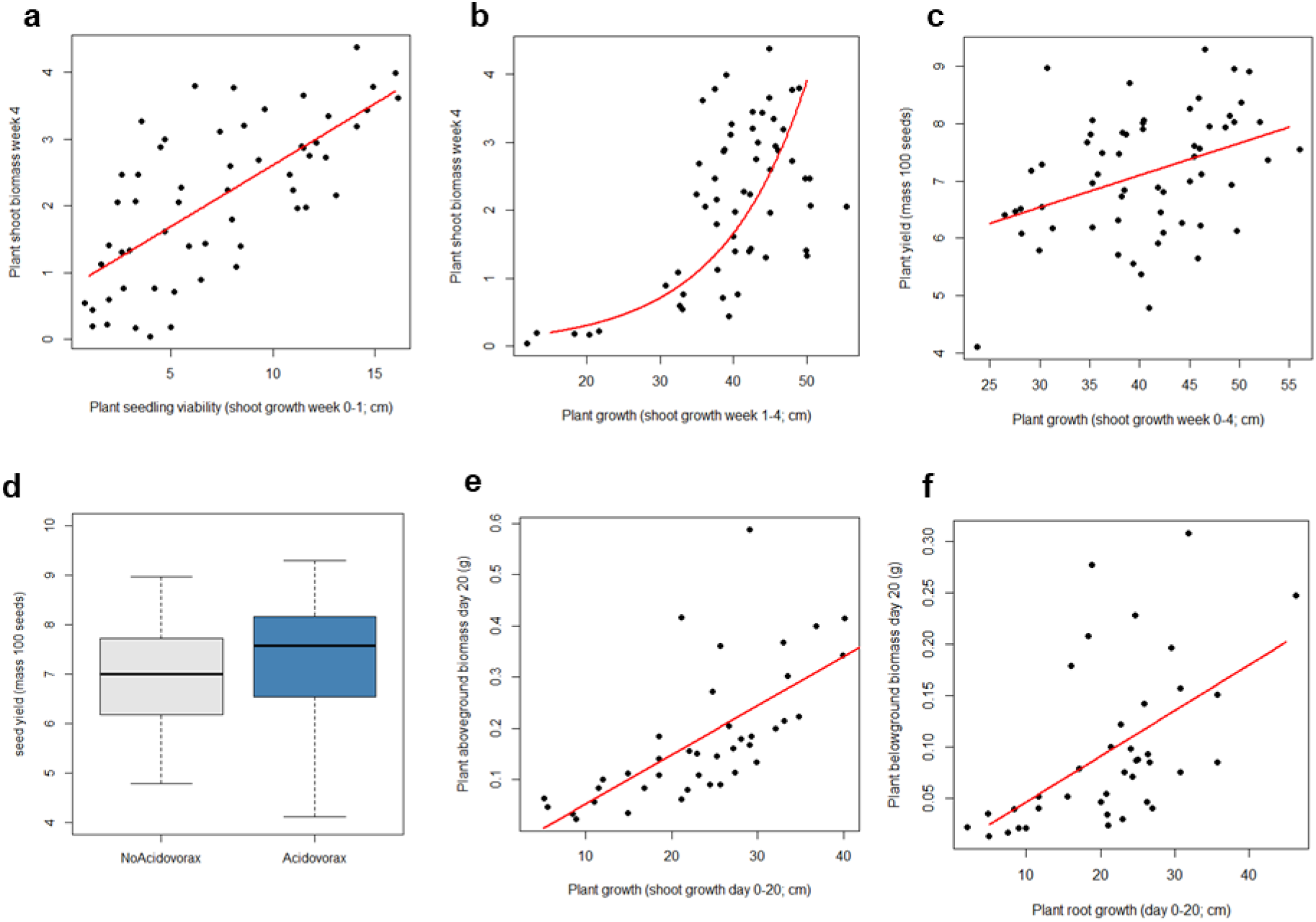
Predictive power of plant shoot length on plant aboveground biomass and yield. Plant growth is measured by plant height difference (longest leaf) from day 0-8, day 8-28, and day 28-56, and as dry total aboveground biomass at day 28 and 56. Plant yield is the mass of 100 seeds at day 84. In a preliminary experiment (unpub. S. Zytynska), (a) early seedling growth (day 0 to day 8) was a good predictor of plant total aboveground biomass at week 4 (day 0-8: F_1,55_=52.30, P<0.001). (b) Aboveground shoot growth from day 8 to day 28 was also a good predictor of plant aboveground biomass (day 8-28: F_1,55_=88.98, P<0.001), however this fitted a logarithmic relationship since plants that grew less than 30 cm in this time produced low biomass (suggesting that there is a threshold of plant height growth for biomass development). This effect changed direction after four weeks of growth (day 28-56), likely associated with changes in resource allocation as the plant begins to produce flowers and seed. (c) Early plant growth (day 0-28) was also associated with final plant yield (mass of 100 seeds) (F_1,58_=10.40, P=0.002), while late plant shoot growth (day 28-56) was not associated with plant yield (F_1,58_=1.14, P=0.289). (d) We also showed that inoculation of *Acidovorax* increased final plant yield (mass of 100 seeds; F_1,50_=5.44, P=0.024). A second experiment (J. Kracht, Master’s thesis TU Munich) confirmed these associations. (e) Plant shoot growth from day 0 until day 20 could predict plant dry biomass at day 20 (F_1,38_=31.81, P<0.001). Further, (f) root length growth and root dry biomass were also highly correlated, indicating a similar predictive power at day 20 for root length to root biomass (F_1,37_=15.77, P<0.001). These results highlight the importance of early plant growth for increased yield, and due to this the current experiment focuses on the effect of the experimental treatments on early growth of the barley plants from germination (day 5-8) to day 22. Plant height difference is used to compare seedling viability (day 5-8, i.e. post-germination), and plant growth (day 8-22).

**Extended data Figure 3.**
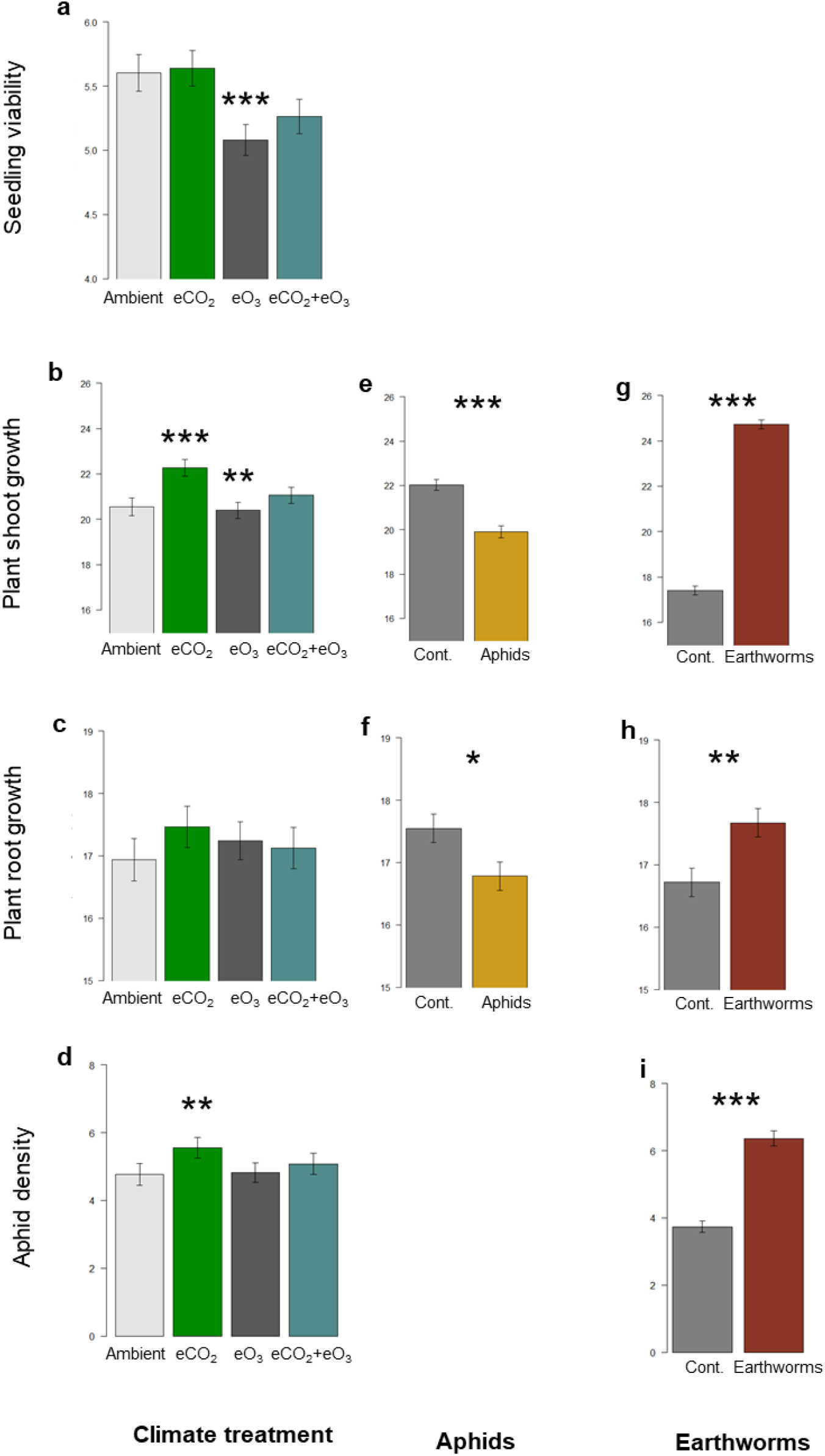
Main effects of abiotic treatment (ambient, elevated CO_2_, elevated O_3_ and combined elevated CO_2_ and O_3_) on plant growth (a-c), *Acidovorax* inoculation (control vs treated) on plant growth (d-f), aphids (absence/presence) on plant growth (h,i), and earthworms (absence/presence) on plant growth (k,l), plus abiotic treatment effects (g) and earthworms (j) on aphid density (aphid number per cm of plant). See Extended Data Table 1 for full results. Error bars +/− 1SE. *** P<0.001, ** P<0.01, * P<0.05

**Extended Data Figure 4.**
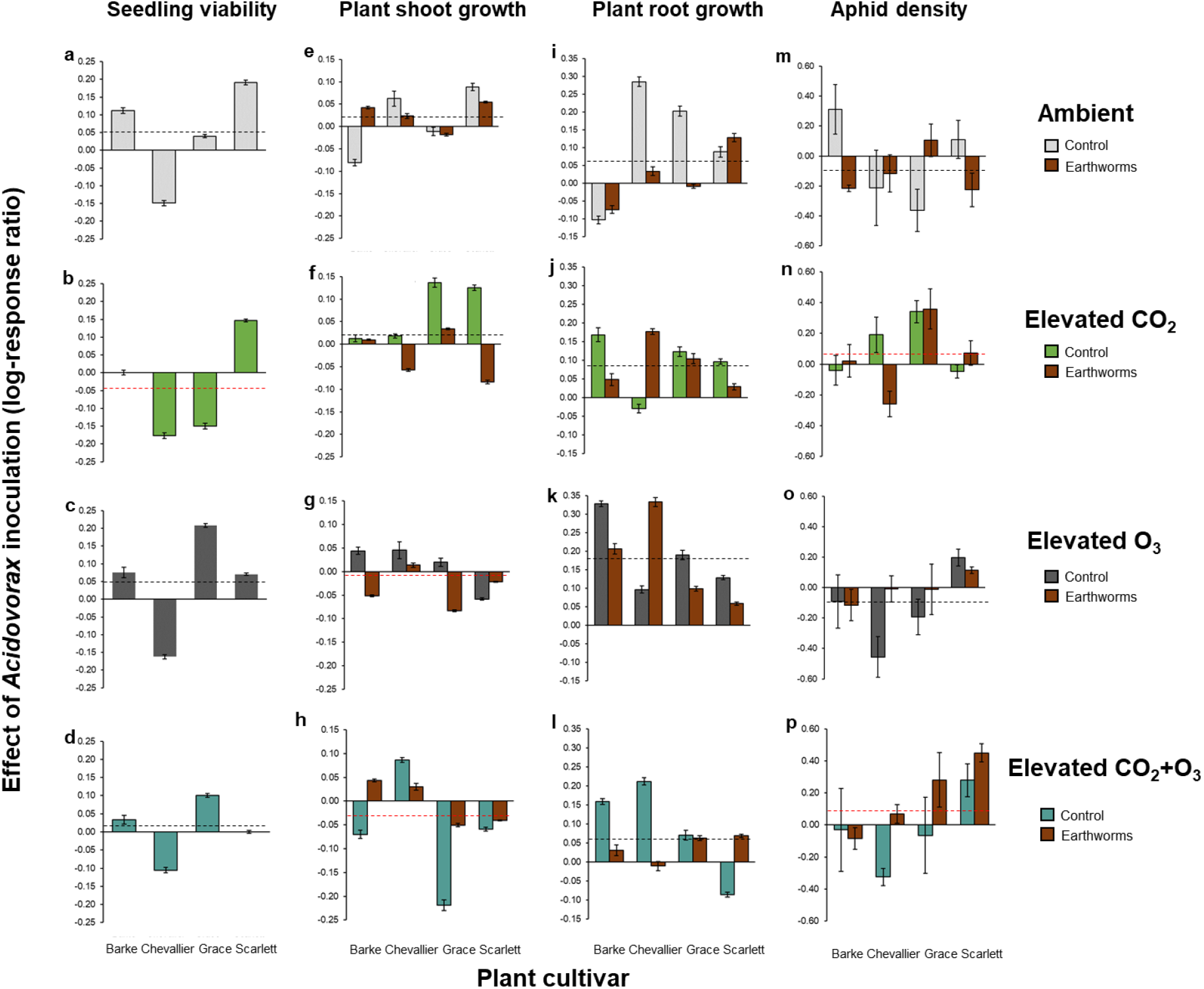
Paired analysis of *Acidovorax* (treated vs control) on plant growth and aphid suppression across abiotic treatments, (a-d) ambient environment, (e-h) elevated CO_2_, (i-l) elevated O_3_, and (m-p) combined elevated CO_2_ and elevated O_3_. Data show effect sizes (log-response ratio) and associated variance estimates (error bars) for each barley cultivar (Barke, Chevalier, Grace, Scarlett) and earthworm treatment (clustered bars, right hand bar is earthworm presence). A positive effect size shows a beneficial effect of Acidovorax on the measured variable, where an effect size of 0.1 represents a response of ~10%. The average effect size across all barley cultivars is shown by the dashed line (black when positive and red when negative for plant health). There was no earthworm treatment for the seedling viability measure.

**Extended Data Figure 5.**
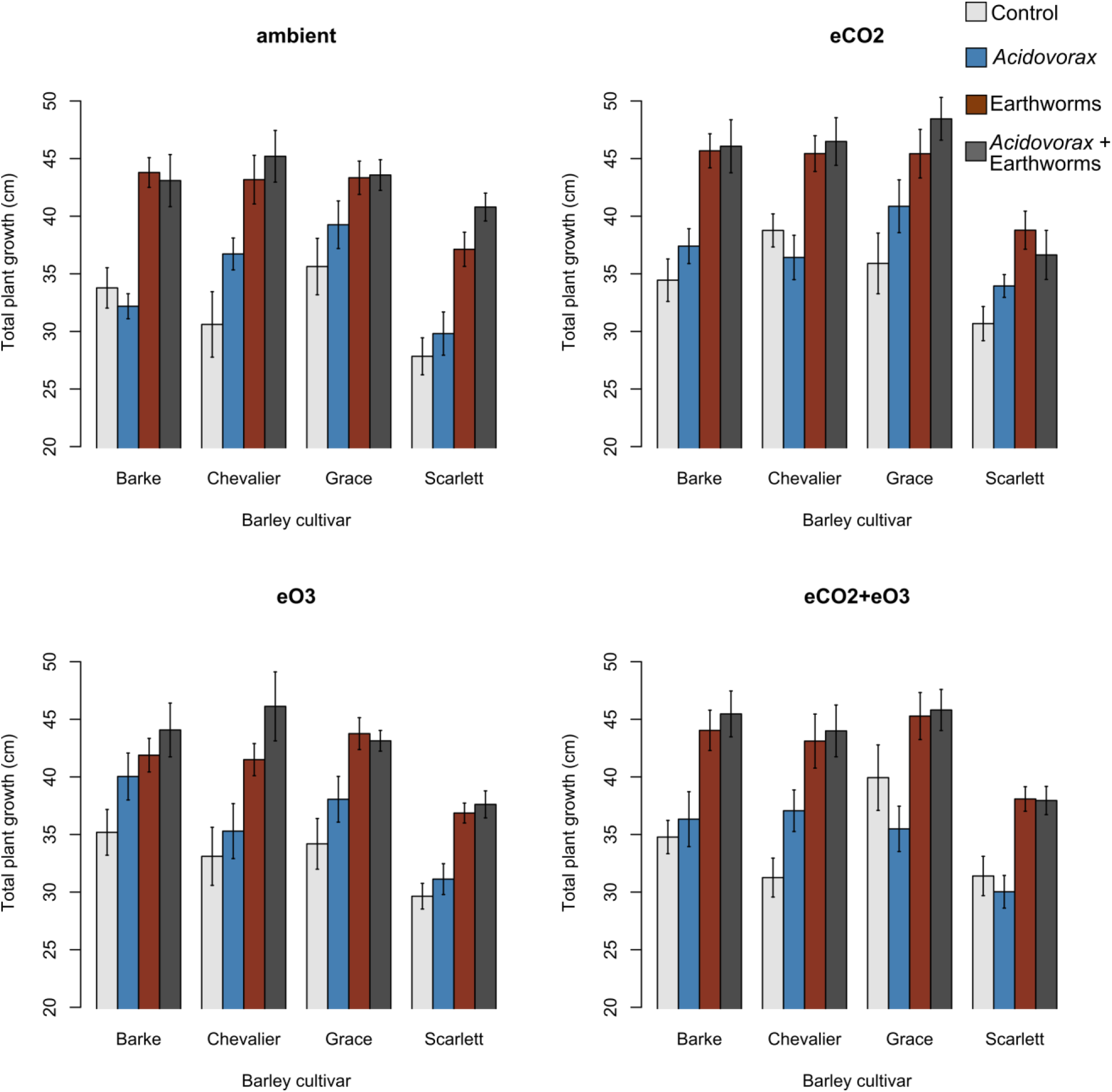
Total plant growth (shoot + root growth) for each barley cultivar within each abiotic environment, across the different *Acidovorax* and earthworm treatments. Error bars +/− 1SE.

**Extended Data Figure 6.**
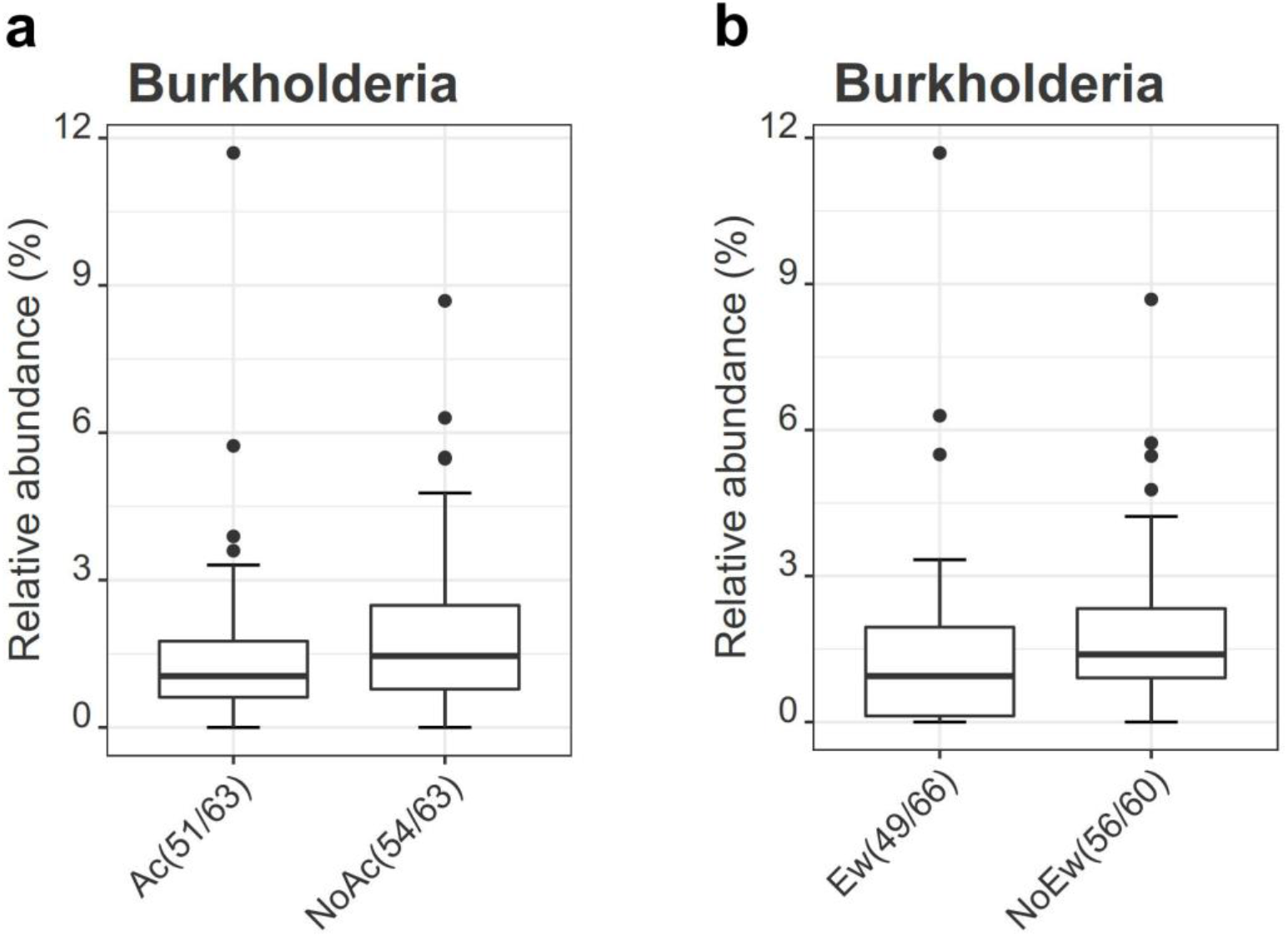
**Serial Group-Comparison depicted as box plots for *Burkholderia* abundance**, which was found to be significantly lower for treatments (a) with *A. radicis* (Ac, p=0.037) and (b) with earthworms (Ew, p=0.012) compared to samples without the respective organisms (NoAc, NoEw). The bottom and top of the box indicate the first and third quartiles, the line inside the box the median and the ends of the whiskers the 10^th^ and 90^th^ percentile values. Outliers are plotted as circles. The second number in brackets indicates how many samples contained sequences allocated to the genus, the first is the number of samples with the respective treatment (i.e. Acidovorax/no Acidovorax, earthworm/no earthworms) adding up to a total of 126 samples. The 4 control samples were not included in this analysis.

