## Extended Data Tables for "Climate change alters beneficial crop-microbe-invertebrate interactions"

### Extended Data Table 1. Summary of linear model results for the plant and aphid response variables

| | | Seedling viability (d5-8) | | | Seedling viability without cv Chevallier | | | Plant growth (d8-d22) | | | Root growth (d5-d22) | | | Aphid load (per cm of plant) | | |
|---|---|---|---|---|---|---|---|---|---|---|---|---|---|---|---|---|
| | df | F | P | | F | P | | F | P | | F | P | | F | P | |
| as.factor(Run) | 2 | 105.15 | <0.001 | *** | 112.83 | <0.001 | *** | 38.46 | <0.001 | *** | 27.51 | <0.001 | *** | 24.01 | <0.001 | *** |
| plant growth variable | 1 | 0.06 | 0.810 | | 0.03 | 0.854 | | 8.25 | 0.004 | ** | 51.19 | <0.001 | *** | na | na | |
| Cultivar | 3 | 90.65 | <0.001 | *** | 112.27 | <0.001 | *** | 55.31 | <0.001 | *** | 11.03 | 0.001 | *** | 2.47 | 0.061 | . |
| Acidovorax | 1 | 1.98 | 0.160 | | 11.17 | <0.001 | *** | 0.41 | 0.521 | | 18.29 | 0.001 | *** | 0.46 | 0.500 | |
| Earthworms | 1 | na | na | | - | - | | 968.66 | <0.001 | *** | 9.98 | 0.002 | ** | 104.29 | <0.001 | *** |
| Aphids | 1 | na | na | | - | - | | 58.27 | <0.001 | *** | 3.73 | 0.054 | . | na | na | |
| eCO2 | 1 | 0.78 | 0.376 | | 0.01 | 0.967 | | 18.35 | <0.001 | *** | 0.03 | 0.861 | | 5.80 | 0.016 | ** |
| eO3 | 1 | 12.86 | <0.001 | *** | 6.35 | 0.012 | * | 5.01 | 0.025 | * | 0.01 | 0.937 | | 0.90 | 0.343 | |
| Cultivar:Acidovorax | 3 | 4.83 | 0.002 | ** | 0.06 | 0.945 | | 0.50 | 0.682 | | 0.45 | 0.719 | | - | - | |
| Cultivar:Aphids | 3 | na | na | | | | | 5.44 | 0.001 | ** | 0.57 | 0.638 | | na | na | |
| Cultivar:Earthworms | 3 | na | na | | | | | 2.70 | 0.045 | * | 0.67 | 0.568 | | - | - | |
| Cultivar:eCO2 | 3 | 1.53 | 0.205 | | 0.86 | 0.423 | | 0.99 | 0.396 | | 1.10 | 0.349 | | 2.54 | 0.036 | * |
| Cultivar:eO3 | 3 | 1.10 | 0.350 | | 0.73 | 0.480 | | 0.84 | 0.474 | | 1.26 | 0.288 | | - | - | |
| Acidovorax:Aphids | 1 | na | na | | | | | 0.46 | 0.498 | | 0.21 | 0.650 | | na | na | |
| Acidovorax:Earthworms | 1 | na | na | | | | | - | - | | 0.48 | 0.489 | | - | - | |
| Acidovorax:eCO2 | 1 | - | - | | 4.71 | 0.030 | * | 0.41 | 0.522 | | 2.65 | 0.104 | | 4.42 | 0.036 | * |
| Acidovorax:eO3 | 1 | 0.09 | 0.770 | | 0.01 | 0.926 | | 3.21 | 0.074 | . | 0.57 | 0.452 | | - | - | |
| Aphids:eCO2 | 1 | na | na | | | | | 0.18 | 0.669 | | 0.18 | 0.668 | | na | na | |
| Aphids:eO3 | 1 | na | na | | | | | 0.39 | 0.531 | | 0.02 | 0.902 | | na | na | |
| Earthworms:Aphids | 1 | na | na | | | | | 0.32 | 0.569 | | 0.32 | 0.572 | | - | - | |
| Earthworms:eCO2 | 1 | na | na | | | | | 2.05 | 0.153 | | 0.02 | 0.900 | | 8.31 | 0.004 | ** |
| Earthworms:eO3 | 1 | na | na | | | | | 2.30 | 0.130 | | 0.01 | 0.925 | | - | - | |
| eCO2:eO3 | 1 | 1.34 | 0.247 | | 2.29 | 0.131 | | 2.88 | 0.090 | . | 0.92 | 0.338 | | - | - | |
| Cultivar:Acidovorax:Earthworms | 3 | na | na | | | | | - | - | | 0.44 | 0.726 | | - | - | |
| Cultivar:Acidovorax:eCO2 | 3 | - | - | | | | | 0.64 | 0.586 | | 0.12 | 0.951 | | - | - | |
| Cultivar:Acidovorax:eO3 | 3 | 3.37 | 0.018 | * | 5.28 | 0.005 | ** | 1.57 | 0.195 | | 0.91 | 0.433 | | - | - | |
| Cultivar:Aphids:eCO2 | 3 | na | na | | | | | - | - | | - | - | | na | na | |
| Cultivar:Aphids:eO3 | 3 | na | na | | | | | 3.96 | 0.008 | ** | - | - | | na | na | |
| Cultivar:Earthworms:Aphids | 3 | na | na | | | | | 5.16 | 0.002 | ** | 2.64 | 0.048 | * | - | - | |
| Cultivar:Earthworms:eCO2 | 3 | na | na | | | | | 2.47 | 0.061 | . | 0.50 | 0.680 | | - | - | |
| Cultivar:Earthworms:eO3 | 3 | na | na | | | | | - | - | | 0.06 | 0.983 | | - | - | |
| Cultivar:eCO2:eO3 | 3 | 3.34 | 0.019 | * | 4.65 | 0.010 | ** | 2.16 | 0.091 | . | 0.01 | 0.999 | | - | - | |
| Acidovorax:Aphids:eCO2 | 1 | na | na | | | | | 0.07 | 0.787 | | 0.96 | 0.329 | | na | na | |
| Acidovorax:Aphids:eO3 | 1 | na | na | | | | | 0.03 | 0.870 | | 2.15 | 0.143 | | na | na | |
| Acidovorax:Earthworms:Aphids | 1 | na | na | | | | | - | - | | 0.05 | 0.822 | | na | na | |
| Acidovorax:Earthworms:eCO2 | 1 | na | na | | | | | - | - | | 0.08 | 0.771 | | - | - | |
| Acidovorax:Earthworms:eO3 | 1 | na | na | | | | | - | - | | 0.00 | 0.995 | | - | - | |
| Acidovorax:eCO2:eO3 | 1 | - | - | | | | | 0.47 | 0.492 | | 1.42 | 0.233 | | - | - | |
| Aphids:eCO2:eO3 | 1 | na | na | | | | | 0.25 | 0.616 | | - | - | | na | na | |
| Earthworms:Aphids:eCO2 | 1 | na | na | | | | | - | - | | 1.31 | 0.253 | | na | na | |
| Earthworms:Aphids:eO3 | 1 | na | na | | | | | 4.59 | 0.032 | * | 2.30 | 0.130 | | na | na | |

| | | | | | | | | | | | | | | | |
|---|---|---|---|---|---|---|---|---|---|---|---|---|---|---|---|
| Earthworms:eCO2:eO3 | 1 | na | na | | | | | - | - | | **3.91** | **0.048** | * | - | - |
| Cultivar:Acidovorax:Earthworms:eCO2 | 3 | na | na | | | | | - | - | | 0.35 | 0.786 | | - | - |
| Cultivar:Acidovorax:Earthworms:eO3 | 3 | na | na | | | | | - | - | | 0.35 | 0.792 | | - | - |
| Cultivar:Acidovorax:eCO2:eO3 | 3 | - | - | | | | | **2.17** | **0.090** | . | 0.95 | 0.416 | | - | - |
| Cultivar:Earthworms:Aphids:eCO2 | 3 | na | na | | | | | - | - | | - | - | | na | na |
| Cultivar:Earthworms:eCO2:eO3 | 3 | na | na | | | | | - | - | | 0.40 | 0.756 | | - | - |
| Acidovorax:Aphids:eCO2:eO3 | 1 | na | na | | | | | **4.02** | **0.045** | * | - | - | | na | na |
| Acidovorax:Earthworms:Aphids:eCO2 | 1 | na | na | | | | | - | - | | 2.03 | 0.154 | | na | na |
| Acidovorax:Earthworms:Aphids:eO3 | 1 | na | na | | | | | - | - | | - | - | | na | na |
| Acidovorax:Earthworms:eCO2:eO3 | 1 | na | na | | | | | - | - | | 0.20 | 0.651 | | - | - |
| Cultivar:Acidovorax:Earthworms:eCO2:eO3 | 3 | na | na | | | | | - | - | | **4.05** | **0.007** | ** | - | - |
| Residual degrees of freedom | | Res df=959 | | | Res df=685 | | | Res df=923 | | | Res df=902 | | | Res df=434 | |
| Weighting factor used to correct for heteroscedasticity | | weighting: varfunc(Cultivar) | | | weighting: varfunc(Cultivar) | | | weighting: varfunc(Cultivar) | | | weighting: varfunc(Cultivar) | | | weighting: varfunc(Cultivar*Acido*Ew) | |

*Values in bold show significant P<0.10, values in grey are kept in the minimal adequate model but not-significant (P>0.10), '-' means that the term did not remain in the minimal adequate model and thus was removed, 'na' means the term was not included in the model (e.g. no aphid or earthworms terms for seedling viability as they were not introduced at that stage).*

*** P<0.001, ** P<0.01, * P<0.05, • P<0.10

**Extended Data Table 2. Summary of matched pairs analysis for effect of *Acidovorax* on plant and aphid response variables**

| **Paired data: *Acidovorax* effect** | | **Seedling viability (d5-8)** | | | **Plant growth (d8-d22)** | | | **Root growth (d5-d22)** | | | **Aphid density** | | |
|---|---|---|---|---|---|---|---|---|---|---|---|---|---|
| | **Df** | **F** | **P** | | **F** | **P** | | **F** | **P** | | **F** | **P** | |
| as.factor(Run) | 2 | - | - | | - | - | | **6.13** | **0.002** | ** | **5.72** | **0.004** | ** |
| Cultivar | 3 | **8.32** | **<0.001** | *** | - | - | | 0.72 | 0.543 | | - | - | |
| Earthworms | 1 | na | na | | 1.23 | 0.267 | | 0.54 | 0.462 | | 0.95 | 0.330 | |
| Aphids | 1 | na | na | | 0.33 | 0.564 | | 0.16 | 0.691 | | na | na | |
| eCO2 | 1 | 3.45 | 0.064 | . | 0.45 | 0.501 | | 2.63 | 0.105 | | **3.98** | **0.047** | * |
| eO3 | 1 | 0.07 | 0.794 | | **4.43** | **0.036** | * | 1.10 | 0.295 | | 1.05 | 0.306 | |
| Cultivar:Earthworms | 3 | na | na | | - | - | | 0.77 | 0.513 | | - | - | |
| Cultivar:eCO2 | 3 | - | - | | - | - | | 0.32 | 0.810 | | - | - | |
| Cultivar:eO3 | 3 | **4.08** | **0.007** | ** | - | - | | 1.81 | 0.145 | | - | - | |
| Aphids:eCO2 | 1 | na | na | | - | - | | **3.11** | **0.078** | . | na | na | |
| Aphids:eO3 | 1 | na | na | | - | - | | **2.74** | **0.099** | . | na | na | |
| Earthworms:Aphids | 1 | na | na | | 2.29 | 0.131 | | - | - | | na | na | |
| Earthworms:eCO2 | 1 | na | na | | 0.23 | 0.628 | | 0.29 | 0.592 | | - | - | |
| Earthworms:eO3 | 1 | na | na | | 1.38 | 0.240 | | 0.02 | 0.881 | | **3.00** | **0.085** | . |
| eCO2:eO3 | 1 | - | - | | 0.02 | 0.894 | | **4.03** | **0.045** | * | - | - | |
| Cultivar:Earthworms:eCO2 | 3 | na | na | | - | - | | 0.38 | 0.766 | | - | - | |
| Cultivar:Earthworms:eO3 | 3 | na | na | | - | - | | 0.21 | 0.893 | | - | - | |
| Cultivar:eCO2:eO3 | 3 | - | - | | - | - | | 1.06 | 0.366 | | - | - | |
| Earthworms:eCO2:eO3 | 1 | na | na | | **5.39** | **0.021** | * | 0.90 | 0.344 | | - | - | |
| Cultivar:Earthworms:eCO2:eO3 | 3 | na | na | | - | - | | **3.90** | **0.009** | ** | - | - | |
| **Residual degrees of freedom** | | Res df=465 | | | Res df=464 | | | Res df=437 | | | Res df=209 | | |
| **Weighting factor used to correct for** heteroscedasticity | | weighting: varfunc(Cultivar) | | | weighting: varfunc(Cultivar) | | | weighting = none | | | weighting = none | | |

*Values in bold show significant P<0.10, values in grey are kept in the minimal adequate model but not-significant (P>0.10), '-' means that the term did not remain in the minimal adequate model and thus was removed, 'na' means the term was not included in the model (e.g. no aphid or earthworms terms for seedling viability as they were not introduced at that stage).*

*** P<0.001, ** P<0.01, * P<0.05, • P<0.10
